# The CARD-CC/Bcl10/paracaspase signaling complex is functionally conserved since the last common ancestor of Planulozoa

**DOI:** 10.1101/046789

**Authors:** Jens Staal, Yasmine Driege, Alice Borghi, Paco Hulpiau, Laurens Lievens, Ismail Sahin Gul, Srividhya Sundararaman, Amanda Gonçalves, Ineke Dhondt, Bart P. Braeckman, Ulrich Technau, Yvan Saeys, Frans van Roy, Rudi Beyaert

## Abstract

Type 1 paracaspases originated in the Ediacaran geological period before the last common ancestor of bilaterians and cnidarians (Planulozoa). Cnidarians have several paralog type 1 paracaspases, type 2 paracaspases, and a homolog of Bcl10. Notably in bilaterians, lineages like nematodes and insects lack Bcl10 whereas other lineages such as vertebrates, hemichordates, annelids and mollusks have a Bcl10 homolog. A survey of invertebrate CARD-coiled-coil (CC) domain homologs of CARMA/CARD9 revealed such homologs only in species with Bcl10, indicating an ancient co-evolution of the entire CARD-CC/Bcl10/MALT1-like paracaspase (CBM) complex. Furthermore, vertebrate-like Syk/Zap70 tyrosine kinase homologs with the ITAM-binding SH2 domain were found in invertebrate organisms with CARD-CC/Bcl10, indicating that this pathway might be the original user of the CBM complex. We also established that the downstream signaling proteins TRAF2 and TRAF6 are functionally conserved in Cnidaria. There also seems to be a correlation where invertebrates with CARD-CC and Bcl10 have type 1 paracaspases which are more similar to the paracaspases found in vertebrates. A proposed evolutionary scenario includes at least two ancestral type 1 paracaspase paralogs in the planulozoan last common ancestor, where at least one paralog usually is dependent on CARD-CC/Bcl10 for its function. Functional analyses of invertebrate type 1 paracaspases and Bcl10 homologs support this scenario and indicate an ancient origin of the CARD-CC/Bcl10/paracaspase signaling complex. Results from cnidarians, nematodes and mice also suggest an ancient neuronal role for the type 1 paracaspases.

## Introduction

The paracaspase MALT1 (PCASP1) was originally identified in humans as an oncogenic fusion with IAP2 in low-grade antibiotic-resistant MALT lymphomas (Dierlamm et al. 1999). Later, it was discovered that MALT1 is a critical component in T and B cell antigen receptor signaling as part of the CARMA1-Bcl10-MALT1 (CBM) signaling complex (Ruefli-Brasse et al. 2003; Ruland et al. 2003; Che et al. 2004). The activated CBM complex aggregates to a filamentous structure (Qiao et al. 2013), where MALT1 subsequently recruits critical downstream proteins, such as TRAF6, for activation of NF-κB-dependent gene expression (Sun et al. 2004) (Figure 1A). CARMA1 belongs to a distinct phylogenetic group which is characterized by a CARD and a coiled-coil (CC) domain and this group of proteins will thus be referred to as CARD-CC proteins. Further studies made it clear that MALT1 plays a role in several different CARD-CC/Bcl10/MALT1 signaling complexes composed of different CARD-CC family proteins: CARD9 (Gross et al. 2006), CARD11 (CARMA1) (Che et al. 2004), CARD14 (CARMA2) (Afonina et al. 2016; Howes et al. 2016; Schmitt et al. 2016) and CARD10 (CARMA3) (McAllister-Lucas et al. 2007). These four CARD-CC proteins are the only members of this family in humans, which makes it unlikely that additional similar CBM complexes will be found. The CARD domain is critical for recruitment of downstream signaling components like Bcl10 (Bertin et al. 2000), whereas the CC domain is critical for oligomerization of the activated signaling complex (Tanner et al. 2007). The use of the different CARD-CC proteins in the CBM complexes is most likely dependent on cell-type specific expression (Scudiero et al. 2013). MALT1 was originally annotated as a “paracaspase” due to sequence similarity with the true caspases and “metacaspases” (Uren et al. 2000). A broader survey of paracaspases in the whole tree of life indicates that “paracaspases” should be considered a sub-class of “metacaspases” that have evolved several times independently (Hulpiau et al. 2016). The name caspase signifies both the structure (cysteine protease) and function (aspartic acid substrate specificity) of the protein family. The semantic association of metacaspases and paracaspases to caspases is therefore unfortunate, since the similar names inspired false assumptions of common roles and properties of the different protein families (Salvesen et al. 2015). There is a stark contrast in the evolution of true caspases and paracaspases. Although both true caspases and paracaspases most likely evolved from a common ancestor in early metazoan evolution, paracaspases have not expanded much and remain remarkably conserved in their domain organization whereas true caspases show an early and rapid expansion, adoption of a wide range of domain organizations and protease substrate specificities (Hulpiau et al. 2016; Moya et al. 2016). Despite the identification of “paracaspase” in 2000, it was not until 2008 that the proteolytic activity of MALT1 was established (Coornaert et al. 2008; Rebeaud et al. 2008). In contrast to true caspases (but similar to metacaspases and orthocaspases), the paracaspase MALT1 cleaves substrates specifically after an arginine residue (Yu et al. 2011; Hachmann et al. 2012; Wiesmann et al. 2012). Lately, some protein substrates have been identified which are cleaved after a lysine by the API2-MALT1 oncogenic fusion (Nie et al. 2015). MALT1 cleaves itself (Baens et al. 2014) and its interacting adaptor protein Bcl10 (Rebeaud et al. 2008), the anti-inflammatory deubiquitinases A20 (Coornaert et al. 2008) and CYLD (Staal et al. 2011), the NF-κB member RelB (Hailfinger et al. 2011), the ubiquitin ligase HOIL-1 (Elton et al. 2015; Klein et al. 2015; Douanne et al. 2016), the specific RNA degradation associated proteins Regnase (Uehata et al. 2013) and Roquin (Jeltsch et al. 2014). The anti-inflammatory role of many of the known protease substrates coupled with the critical role for MALT1 in pro-inflammatory signaling has sparked an interest in targeting MALT1 protease activity as a therapeutic strategy treatment of autoimmune diseases (Mc Guire et al. 2014). The proteolytic activity of MALT1 was also found to be critical for certain cancers (Juilland and Thome 2016), which has sparked an interest in MALT1 protease activity as a cancer therapy target. Although the MALT1 scaffold function for recruitment of downstream TRAF6 has been clearly associated to NF-κB activation (Noels et al. 2007), the MALT1 protease activity plays a more subtle role being specifically required for c-Rel activation (Ferch et al. 2007; Gringhuis et al. 2011; Hailfinger et al. 2011; Baens et al. 2014). There is some evidence that MALT1 also regulates or cross-talks with other pathways, such as JNK/AP-1 (Staal et al. 2011), mTORC1 (Hamilton et al. 2014), RhoA/ROCK (Klei et al. 2016), MYC (Dai et al. 2016) and possibly WNT (Bognar et al. 2016). MALT1 belongs to the type 1 paracaspase family, which consists of an N-terminal death domain, immunoglobulin domains and a paracaspase domain (Hulpiau et al. 2016). The type 1 family of paracaspases originated sometime during the Ediacaran geological period, preceding the last common ancestor of bilaterians and cnidarians (Peterson et al. 2004; Knoll et al. 2006; Hulpiau et al. 2016). The cnidarians (e.g. jellyfish, sea anemone, hydra and coral) and bilaterians (e.g. vertebrates, insects, nematodes, mollusks and ringed worms) form the planulozoan clade (Dunn et al. 2014). In our previous survey of paracaspases and MALT1-associated proteins, type 1 paracaspases and Bcl10 could not be found outside planulozoa (Hulpiau et al. 2016). Cnidarians typically contain several paralogs of both type 1 and the ancient type 2 paracaspases whereas bilaterians typically contain a single copy of a type 1 paracaspase. A notable exception is the jawed vertebrates, where the type 1 paracaspase got triplicated. Subsequently, two paralogs were lost in the mammalian lineage leaving PCASP1 (MALT1) as the single paracaspase in mammals (Hulpiau et al. 2016). Importantly, some organisms, such as the nematode *Caenorhabditis elegans*, contain a conserved type 1 paracaspase but lack NF-κB (Sullivan et al. 2009), which indicate that other roles or mechanisms might be responsible for the conservation of the general domain organization of the type 1 paracaspases (Hulpiau et al. 2016). The WormBase *C. elegans* phenotype database indicates an important role for the type 1 paracaspase (*tm289* vs *tm321* mutant) in nematodes where loss of paracaspase causes a “sterile or lethal” phenotype (C. elegans Deletion Mutant Consortium 2012). On the other hand, despite the apparent importance and remarkable conservation of type 1 paracaspases, there are examples of bilaterians that have lost their paracaspase - most notably the group of flies including the fruit fly *Drosophila melanogaster* (Hulpiau et al. 2016). This indicates that alternative mechanisms can take over the unknown role which is usually filled by the type 1 paracaspases in most other planulozoan organisms. Mice deficient for MALT1 do not show any obvious developmental phenotypes, and primarily show defects in T and B cell functions (Ruefli-Brasse et al. 2003; Ruland et al. 2003). The T and B cell antigen receptor signaling pathways are currently the most extensively investigated pathways for MALT1 activity (Demeyer et al. 2016). Humans deficient for MALT1 show similar symptoms which manifest as potentially lethal immunodeficiency. Treatment of MALT1 deficient patients with healthy donor adaptive immune cells results in a remarkable recovery (Punwani et al. 2015; Rozmus et al. 2016), supporting the idea that the most acute effects of MALT1 deficiency in humans and other mammals are related to the adaptive immune system. Apart from functional studies of MALT1 in human cells, patients and mouse models - investigating the evolutionary history of the type 1 paracaspases, their interacting proteins and molecular functions in alternative model systems could provide important clues to yet-unknown roles and functions of human MALT1 (Hulpiau et al. 2016). Finding those alternative functions of MALT1 could also ultimately be important for future MALT1 inhibitor-based therapies (Demeyer et al. 2016) and to identify yet undiscovered issues that might affect MALT1 deficient patients.

**Figure 1:**
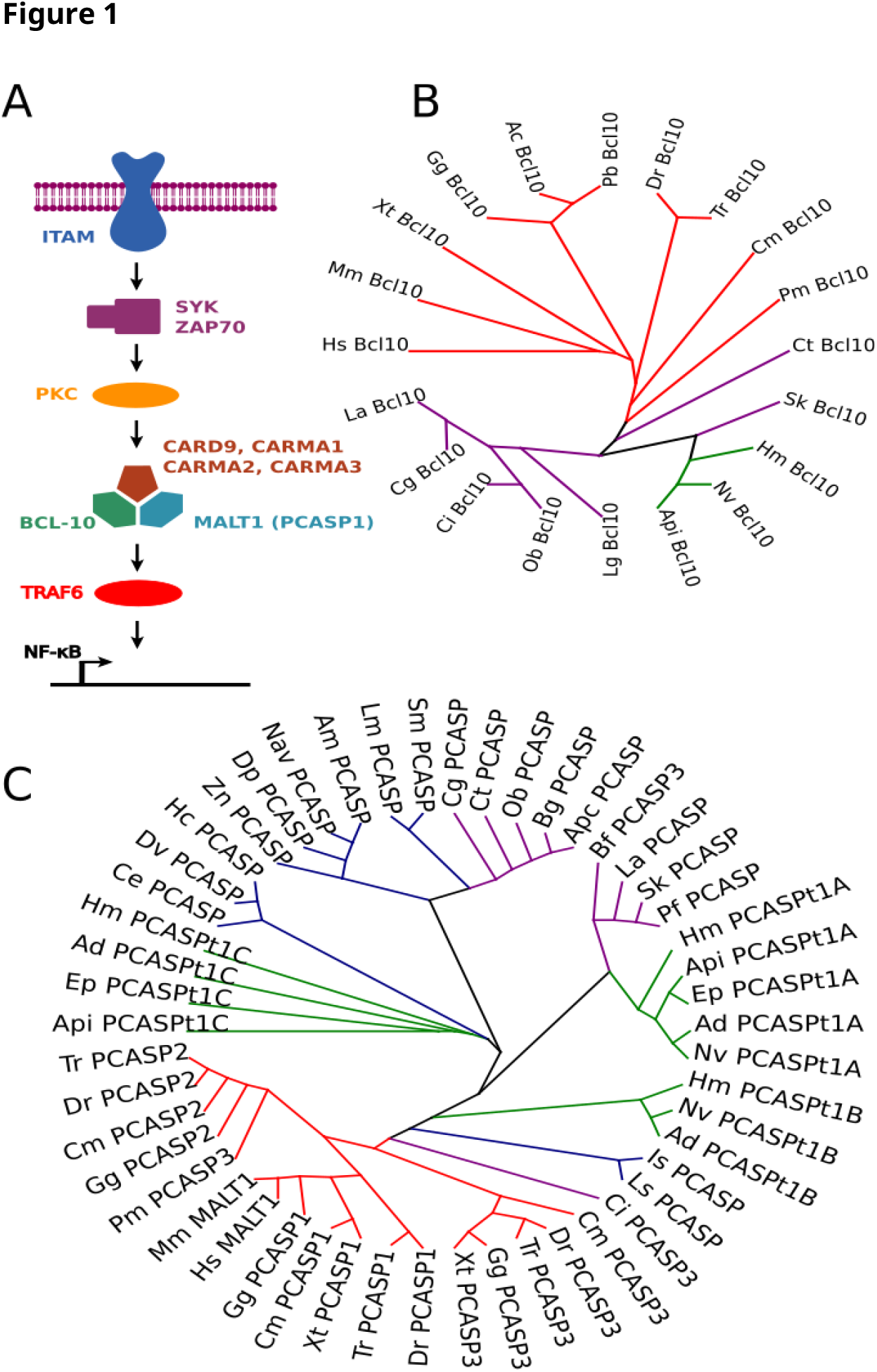
Phylogeny of Bcl10 and type 1 paracaspase N-terminal sequence. **A)** A simplified overview of CBM complex-mediated activation of NF-κB signaling in human and mouse. **B)** Maximum likelyhood phylogeny (MUSCLE + PhyML) of Bcl10. **C)** Maximum likelihood phylogeny (MUSCLE + PhyML) of the type 1 paracaspase DD-Ig1-Ig2 N-terminal domain, which is likely to be involved in Bcl10-binding. Species key: <u>Vertebrates</u>: Hs=Human, Mm=mouse, Gg=Chicken, Xt=African clawed frog, Dr=Zebrafish, Tr=Fugu, Cm=Elephant shark, Pm=Sea lamprey. <u>Tunicates</u>: Ci=Vase tunicate <u>Lancelets</u>: Bf=Florida lancelet. <u>Hemichordates</u>: Sk=Acorn worm Pf=Hawaiian acorn worm <u>mollusks</u>: Cg=Pacific oyster, Lg=Limpet, Ob=Califonia two-spot octopus <u>Brachiopods</u>: La=*L. anatina* <u>Annelids</u>: Ct=polychaete worm <u>Arthropods</u>: Am=Honey bee, Nav=jewel wasp, Sm=African social velvet spider, Pt=common house spider, Is=Fire ant, Lm=Horseshoe crab, Zn=termite <u>Nematodes</u>: Ce, Dr, Hc <u>Cnidaria</u>: Nv=Starlet sea anemone, Ep=sea anemone, Hm=Hydra, Ad=Stag horn coral. Vertebrates highlighted in red branches, bilaterian invertebrate species with Bcl10 with purple branches, cnidaria in green and species from phyla completely lacking Bcl10 (e.g. arthropods, nematodes) in blue.

## Results & Discussion

### Correlation vertebrate-like type 1 paracaspases and presence of Bcl10

While searching for invertebrate homologs of type 1 paracaspases and Bcl10, it became apparent that type 1 paracaspases from species containing Bcl10 generally had higher BLASTp rankings compared to species from phyla lacking *Bcl10*. Bcl10 sequences in vertebrates appear to evolve in a manner similar to how the species have diverged throughout evolution, while the invertebrate Bcl10 sequences are poorly resolved (Figure 1B). To get a better understanding of early Bcl10 evolution, more sequences from invertebrate genomes are needed (GIGA community of scientists 2014; Long et al. 2016). Different alignment strategies and phylogenetic analyses of several type 1 paracaspases verify that type 1 paracaspases from bilaterian species that contain *Bcl10* (mollusks, annelids, hemichordates) often cluster closer to the vertebrate paracaspases, either directly or indirectly by clustering with the invertebrate *Pcasp3* orthologs from tunicate and lancelet (Hulpiau et al. 2016) (Figure S1), indicating a conserved common Bcl10-dependent ancestor. The phylogenetic topology of the type 1 paracaspases is strongly dependent on the sequence segments used for the analysis, indicating that different parts of the protein have experienced different types of selection pressure. Since the N-terminal death domain and immunoglobulin domain are associated to Bcl10 binding (Langel et al. 2008), a phylogenetic analysis of the sequence N-terminal of the caspase-like domain was performed. This analysis shows a stronger association between paracaspases from *Bcl10*-containing species, but does not cluster known paralogs within the deuterostomes (Figure 1C). We can currently not resolve whether there were two paracaspase paralogs, one Bcl10-dependent and the other Bcl10-independent already from the planulozoan last common ancestor or if Bcl10-independent paralogs have evolved several times. According to some models, the cnidarian paracaspases tend to cluster together, which would indicate that the last common planulozoan ancestor had a single Bcl10-dependent type 1 paracaspase which later expanded independently in the bilaterian and cnidarian lineages. Phylogenetic analyses of full-length paracaspases and of the highly conserved caspase-like domain, however, indicate that the last common bilaterian ancestor had 2-3 different type 1 paracaspases (Figure S1). Since nematode paracaspase sequences tend to cluster closer to mollusk sequences rather than arthropod sequences, it is possible that Bcl10 was lost independently in the arthropod and nematode lineages. Because of the unclear early bilaterian evolutionary history of the type 1 paracaspases, only deuterostome paracaspases, which are clear orthologs of the vertebrate *Pcasp3*, are currently classified and named as *Pcasp3*. The three cnidarian type 1 paracaspase paralogs are currently annotated “A” to “C” in order to avoid possible future name space conflicts when the invertebrate paracaspases can be more accurately classified and numbered.

### Functional conservation of invertebrate type 1 paracaspases

Based on BLASTp and subsequent phylogenetic analyses, the mollusk paracaspases were identified as the non-deuterostome homologs most closely resembling vertebrate type 1 paracaspases (Hulpiau et al. 2016). The pacific sea oyster (*Crassostrea gigas*) (Zhang et al. 2012) was selected as a model and cDNA source (Fleury et al. 2009) for the mollusks. Conversely, the most distantly related species where type 1 paracaspases and *Bcl10* could be found are cnidaria (Hulpiau et al. 2016). The cnidarian model organism starlet sea anemone (*Nematostella vectensis*) (Putnam et al. 2007) was used as a cDNA source for as divergent homologous proteins as possible. To investigate the functional conservation of invertebrate type 1 paracaspases, we transiently transfected MALT1-deficient HEK293T cells with several type 1 paracaspases fused to the *ETV6* HLH domain, leading to their oligomerization and artificial activation (Baens et al. 2014). Transfected activated paracaspases were analyzed for scaffold function by activation of NF-κB dependent luciferase reporter expression and for protease function by CYLD cleavage. As positive control, the currently most distantly related vertebrate paracaspase with conserved activity (zebrafish PCASP3) (Hulpiau et al. 2016) was used. In an NF-κB dependent luciferase reporter assay, only the activated zebrafish PCASP3 could induce the reporter to relevant levels, indicating that the pacific oyster (CgPCASP) and the two starlet sea anemone type 1 paracaspase paralogs (NvPCASP-t1A, NvPCASP-t1B) could not recruit and activate critical downstream signaling components (Figure 2A). Although a statistically significant NF-κB induction could be seen from CgPCASP, the levels were more than 150-fold less than what is observed from vertebrate paracaspases and probably not relevant (Figure 2A). CYLD is chosen as model substrate for evaluation of paracaspase protease activity and specificity since it is a large protein with many potential aspecific cleavage sites, and it represents one of the oldest paracaspase substrates (Hulpiau et al. 2016). Exceptionally, the sea anemone (*Nematostella* and *Aiptasia*) CYLD homolog is short and roughly corresponds to the C-terminal cleavage fragment of human CYLD. Other cnidarians (e.g. the coral *Acropora digitifera* and the hydrozoan *Hydra vulgaris*) show a long form of CYLD which aligns to the full-length sequence of human CYLD. Evaluation of protease activity on the human CYLD substrate revealed that the pacific oyster paracaspase specifically cleaves human CYLD at R324, just like vertebrate paracaspases (Figure 2B). This differs from our previous studies of invertebrate paracaspases such as the type 1 paracaspase from *C. elegans* and the more distantly related type 2 paracaspases, which failed to show any activity (Hulpiau et al. 2016). On the other hand, the “A” and “B” type 1 paracaspase paralogs from *N.vectensis* could not cleave CYLD at all, indicating that paracaspase substrate specificity is not conserved in the cnidarians despite being an organism with a *Bcl10* homolog. It is however important to stress that a lack of MALT1-like activity of a distant homolog in a human host cell does not exclude MALT1-like activity in its native cellular environment. Many critical parameters might differ between the cellular environments such as interacting host proteins, post-translational modifications (*e.g*. phosphorylation, ubiquitination) and biophysical conditions (*e.g*. temperature, pH, redox state, ion concentrations). Previous studies with heterologous expression of cnidarian caspase-8 homologs have however been able to establish functional conservation in a human cellular background (Sakamaki et al. 2014). Nevertheless, we can establish that the MALT1-like protease substrate specificity predates the divergence of deuterostomian and protostomian bilaterians and that the MALT1-like protease substrate specificity probably predate the MALT1-like scaffold function for induction of NF-κB.

**Figure 2:**
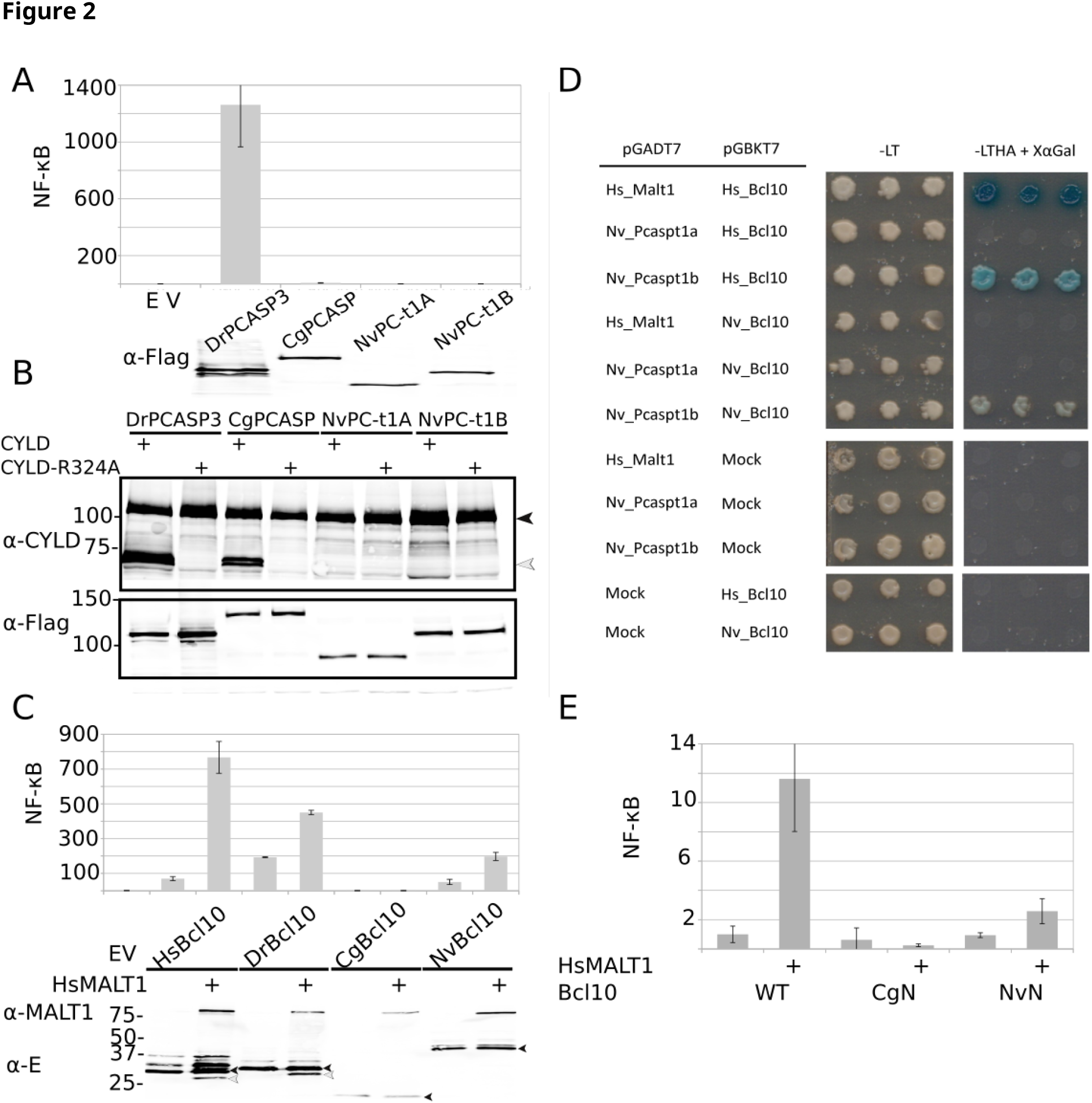
Functional conservation of invertebrate paracaspase and Bcl10. **A)** NF-κB-dependent luciferase reporter gene induction by activated HLH-paracaspase fusion proteins expressed in MALT1 deficient HEK293T cells together with an NF-κB-dependent luciferase reporter expression plasmid and a constituitively expressed β-galactosidase reporter gene. Luciferase values are normalized against β-galactosidase and expressed as fold induction compared to samples not expressing a HLH-paracaspase fusion (EV). Error bars represent 95% confidence intervals (Student’s t-distribution). The lower part of the panel shows the expression of each Flag-tagged HLH-fused type 1 paracaspase as revealed by western blotting and development with anti-Flag. Experiments were repeated at least twice. **B)** CYLD cleavage by activated HLH-paracaspase fusion proteins. MALT1 deficient HEK293T cells were transiently transfected with the indicated HLH-paracaspases together with either human WT CYLD or CYLD(R324) in which the MALT1 cleavage site is mutated. Cell lysates were analyzed for CYLD expression and cleavage via western blotting and detection with anti-CYLD. Human CYLD is specifically cleaved by vertebrate paracaspases after residue R324, resulting in a 70kDa C-terminal fragment. Cleavage of WT CYLD but failure to cleave the R324A mutant indicates a conserved substrate specificity. To show equal expression of each HLH-fused type 1 paracaspase, the blot was also developed with anti-Flag (lower part). Experiments were repeated at least twice. **C)** Human MALT1-dependent NF-κB induction by different Bcl10 homologs. The indicated Bcl10 homologs were expressed in MALT1 deficient HEK293T cells together with an NF-κB-dependent luciferase reporter expression plasmid and a constituitively expressed β-galactosidase reporter gene. Where indicated human MALT1 was also co-expressed. Luciferase values are normalized against β-galactosidase and expressed as fold induction compared to samples not expressing Bcl10 (EV). Error bars represent 95% confidence intervals (Student’s t-distribution). Bcl10 induces NF-κB via MALT1, which is illustrated by the increase of luciferase activity when the cells are reconstituted with human MALT1. Expression of E-tagged Bcl10 homologs and human MALT1 were revealed by western blotting and detection with anti-E-tag or anti-MALT1, respectively (lower part). Full-length Bcl10 is indicated by black arrow heads and cleaved Bcl10 by white arrow heads. Experiments were repeated at least twice.**D)**Yeast-2-hybrid assay demonstrating conserved interaction between *N. vectensis* and human Bcl10 and type 1 paracaspases. Left panel represents growth on non-selective media and right panel (-LTHA) selective growth, where only a combination of bait and prey clones with interacting proteins can grow. As an independent readout, strong interactions also stain blue from X-gal. **E)** Bcl10 CARD-dependent MALT1-mediated NF-κB induction. Wild-type human Bcl10 and hybrid Bcl10 clones where residues 1-102 in human Bcl10 is replaced by the corresponding residues from *C.gigas* (CgN) or *N. vectensis* (NvN)were expressed in MALT1 deficient HEK293T cells together with an NF-κB-dependent luciferase reporter expression plasmid and a constituitively expressed β-galactosidase reporter gene. Where indicated human MALT1 was also co-expressed. Luciferase values are normalized against β-galactosidase and expressed as fold induction compared to samples expressing human wild-type Bcl10 without MALT1. Experiments were repeated at least twice.

### Functional conservation of Bcl10-induced MALT1 activity

To further investigate the functional conservation of the Bcl10/paracaspase co-evolution, we transfected human, zebrafish, pacific oyster and starlet sea anemone Bcl10 in MALT1 deficient HEK293T cells with or without reconstitution with human MALT1. Strikingly, the starlet sea anemone Bcl10 could induce human MALT1-mediated NF-κB induction. This result is highly unexpected, since a critical MALT1 Ig domain interaction sequence (residues 107-119 in human Bcl10) that has been identified downstream of the CARD domain in human Bcl10 (Langel et al. 2008) can only be found in vertebrates. Importantly, the critical 107-119 residues in human Bcl10 are not conserved in zebrafish, demonstrating that alternative additional C-terminal paracaspase binding domains are common. In contrast to human and zebrafish Bcl10, NvBcl10 does not appear to be cleaved by human MALT1 (Figure 2C). The observation that cnidarian Bcl10 can activate human MALT1 indicates a highly conserved interaction surface between the two proteins. A conserved Bcl10-paracaspase interaction was confirmed with yeast-2-hybrid analysis, where the *N. vectensis* type 1 paracaspase “B” paralog readily interacted with both human and *N. vectensis* Bcl10 (Figure 2D). The *N. vectensis* type 1 paracaspase “A” paralog did not show any interaction with Bcl10. Interestingly, this difference in Bcl10 interaction is reflected by the phylogenetic analysis of the N-terminal sequence of type 1 paracaspases, where the cnidarian “B” paralog clusters closer to type 1 paracaspases from vertebrates and Bcl10-containing invertebrate bilaterian species (Figure 1C). In contrast to the functional interaction revealed in the NF-κB luciferase assays (Figure 2C), no physical interaction could be established between *N. vectensis* Bcl10 and human MALT1 by yeast-2-hybrid (Figure 2D) or co-immunoprecipitation (not shown). An ancient Bcl10/paracaspase interaction is highly interesting, since a conserved protein-protein interaction could be used to further model the critical interaction surfaces using evolutionary data (Hopf et al. 2014). The pacific oyster Bcl10 failed to induce any NF-κB reporter activity. The currently annotated pacific oyster and other mollusk Bcl10 homologs are smaller than Bcl10, and only consist of a clear alignment of the CARD domain up until residue 102 of human Bcl10. This N-terminal region has been shown to be required but insufficient for NF-κB induction (Langel et al. 2008). Importantly, the acidic residues required for interaction with upstream CARD-CC proteins (Li et al. 2012) are conserved as far back as *N. vectensis*. In addition to the 102 conserved N-terminal residues, all Bcl10 homologs show a proline-rich motif close to the C-terminus. The functional importance of this proline-rich region is however unknown, since it is dispensable for MALT1 activation. For further proof that the Bcl10 CARD domain is functionally conserved, we generated hybrid Bcl10 clones where residues 1-102 in human Bcl10 are replaced by the corresponding residues from *N. vectensis* or *C. gigas* Bcl10. In addition we generated the minimal active human Bcl10 1-119 construct and also replaced residues 1-102 in this construct with sequences from *N. vectensis* and *C. gigas*. These hybrid Bcl10 clones showed the same MALT1-dependent NF-κB activation pattern as the wild-type clones (Figure 2C,E). Since the pacific oyster hybrid Bcl10 still failed to induce human MALT1 activity, we can conclude that the CARD domain-dependent paracaspase activation has undergone a lineage-specific divergence. In contrast, *N. vectensis* hybrid Bcl10 shows the same relative level of induction as the wild-type *N. vectensis* Bcl10, indicating that the sub-optimal induction is due to the CARD domain and not due to missing interaction domains at the C-terminal part of Bcl10. Although sub-optimal (2.5-fold vs 10-fold MALT1-dependent NF-κB induction), we have demonstrated that the CARD domain, which is critical for paracaspase interaction, is conserved despite very low sequence identity in the corresponding sequence from *N. vectensis* (35% identity) whereas the closer related *C. gigas* (40% identity) lacks the ability activate human MALT1. Both distant homologs have acidic residues that correspond to E84, a glutamine corresponding to Q92 and hydrophobic residues corresponding to residues L95, I96 & I99 in human Bcl10, which all are critical for MALT1 interaction (Langel et al. 2008). On the other hand, only *Nematostella* have an acidic residue that correspond (but shifted one position) to the critical D80 residue in human Bcl10. From these experiments we can conclude that the Bcl10/paracaspase interaction is likely to be ancient and that the critical Bcl10 CARD/paracaspase interaction is highly conserved. We also demonstrate that the non-CARD component of Bcl10 required for MALT1 activation is not dependent on a single highly conserved MALT1-binding peptide sequence downstream of the CARD domain.

### Cnidarian-level functional conservation of downstream signaling proteins

Since neither mollusk nor sea anemone type 1 paracaspases were able to induce NF-κB in a human cellular background, we wanted to investigate whether downstream signaling components are functionally conserved. The TRAF family of E3 ubiquitin ligases are conserved and diverged before the cnidarian/bilaterian last common ancestor (Meyer and Weis 2012). In humans, TRAF6 is the critical member of this family for signaling downstream of MALT1 (Sun et al. 2004). In other signaling pathways or in the API2-MALT1 oncogenic fusion, TRAF2 plays an as important role in NF-κB induction (Noels et al. 2007; Borghi et al. 2016). The Ig2 TRAF6 binding motif (TDEAVECTE) and the C-terminal (PVETTD) TRAF6-binding site in MALT1 (Noels et al. 2007) are PCASP1-specific, but we know that vertebrate PCASP2 and PCASP3 paralogs still are as efficient in NF-κB induction (Hulpiau et al. 2016). One TRAF6 binding (TPEETG) site in human MALT1 appears to be conserved in all vertebrate paralogs, and the corresponding critical glutamic acid might be present in mollusk and cnidarian paracaspases while it appears to be missing in the nematode and arthropod homologs. To investigate whether the type 1 paracaspase - TRAF interaction has undergone lineage-specific divergence, we cloned the *N. vectensis* homologs of *Traf2* and *Traf6* and co-expressed them with the two *N. vectensis* type 1 paracaspase paralogs fused to the activating *ETV6* HLH domain in HEK293T cells to test them in an NF-κB luciferase reporter assay (Figure 3). The cnidarian TRAF2 and TRAF6 homolog proteins were both highly efficient in inducing NF-κB in a human cellular background. In contrast to what would have been expected if a *N. vectensis* type 1 paracaspase would have recruited and activated one of the *N. vectensis* TRAF homologs, no synergistic induction of NF-κB could be seen. This indicates that the evolution of type 1 paracaspases as NF-κB inducing scaffold proteins by recruitment and activation of TRAF6 occurred later, possibly after the deuterostomian / protostomian split in the bilaterian lineage.

**Figure 3:**
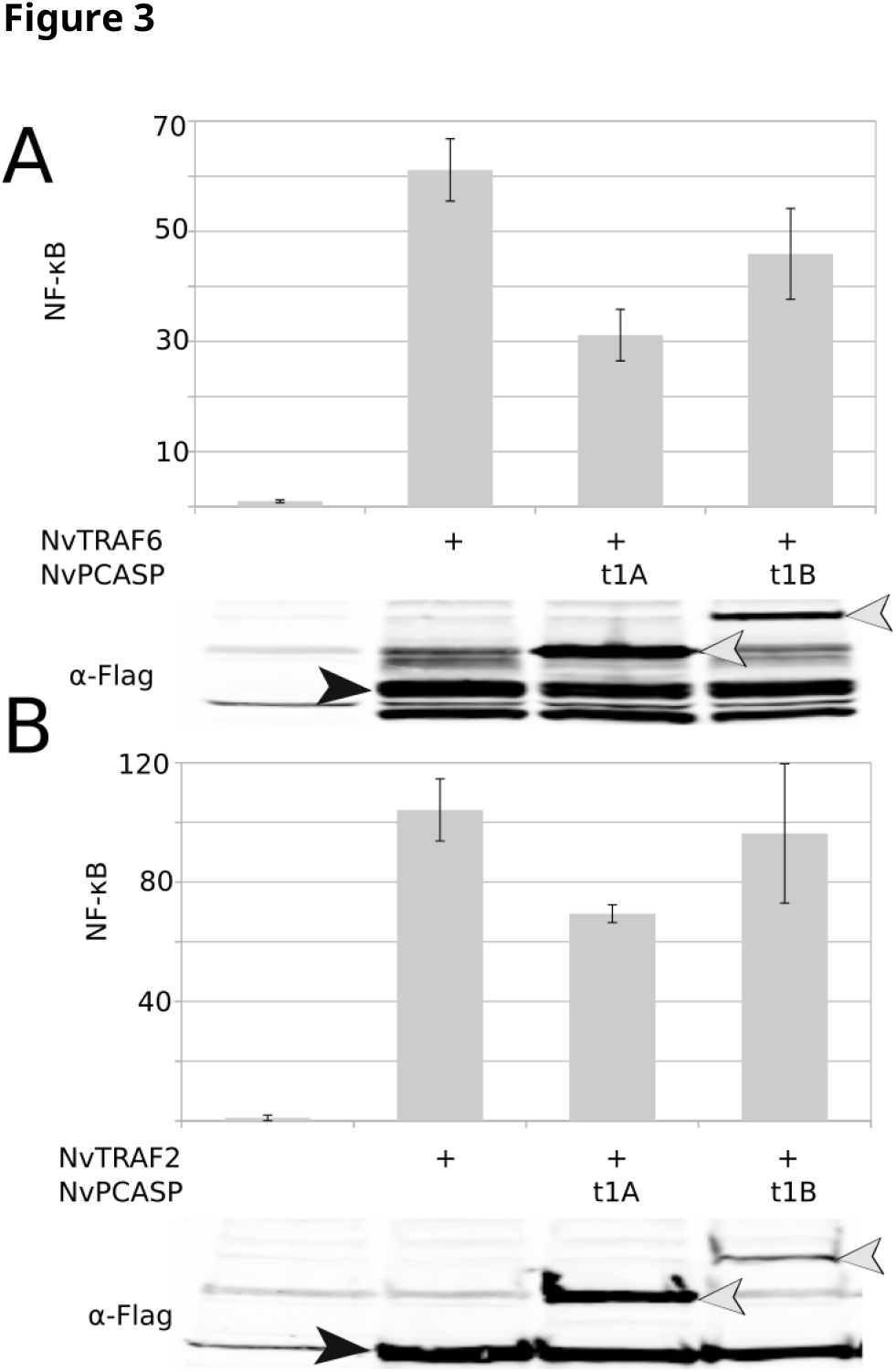
Functional conservation of cnidarian TRAF6 and TRAF2. **A)** *N. vectensis* TRAF6 was expressed in human HEK293T cells NF-κB-dependent luciferase reporter expression plasmid and a constituitively expressed β-galactosidase reporter gene. TRAF6 was transfected either alone or together with the indicated *N. vectensis* HLH-paracaspase fusion protein. Luciferase values are normalized against β-galactosidase and expressed as fold induction compared to samples not expressing TRAF6. Error bars represent 95% confidence intervals (Student’s t-distribution). The lower part of the panel shows the expression of TRAF6 (black arrow head) and each Flag-tagged HLH-fused type 1 paracaspase (white arrow heads) as revealed by western blotting and development with anti-Flag. Experiments were repeated at least twice. **B)** *N. vectensis* TRAF2 was expressed in human HEK293T cells NF-κB-dependent luciferase reporter expression plasmid and a constituitively expressed β-galactosidase reporter gene. TRAF2 was transfected either alone or together with the indicated *N. vectensis* HLH-paracaspase fusion protein. Luciferase values are normalized against β-galactosidase and expressed as fold induction compared to samples not expressing TRAF2. Error bars represent 95% confidence intervals (Student’s t-distribution). The lower part of the panel shows the expression of TRAF2 (black arrow head) and each Flag-tagged HLH-fused type 1 paracaspase (white arrow heads) as revealed by western blotting and development with anti-Flag. Experiments were repeated at least twice.

### Conservation and co-evolution of the CBM complex

Previous studies have shown that the MALT1-like protease and scaffold activities are conserved at least as far back as the last common ancestor of the three jawed vertebrate type 1 paracaspase paralogs (Hulpiau et al. 2016). Similarly, also Bcl10 has been shown to be functionally conserved as far back as zebrafish, as is the upstream interaction to CARMA proteins (Mazzone et al. 2015). We have now shown that Bcl10 and MALT1-like activities from type 1 paracaspases are considerably older (Figure 2), most likely preceding the Cambrian explosion (Dunn et al. 2014). The observation that invertebrate type 1 paracaspases from organisms that also contain Bcl10 are more similar to the vertebrate paracaspases (Figure 1C) provides a new interesting perspective on the functional evolution of MALT1. CARMA proteins are unique to jawed vertebrates, but the conserved CARD-CC domain organization can be found in some invertebrates. A likely evolutionary scenario for the CARMA proteins is that a CARD9-like CARD-CC got fused with a ZO-1/Dlg5-like MAGUK protein upstream of the PDZ domain early in the jawed vertebrate evolution (de Mendoza et al. 2010). The placement of CARD14 (CARMA2) at the base of the CARMA/CARD9 proteins found in vertebrates based on the CARD domain phylogeny (Figure 4A) is consistent with phylogenies made with the MAGUK domain (de Mendoza et al. 2010), indicating that CARD14 might be the ancestral or least diverged CARMA in vertebrates. Interestingly, the presence of three CARMA paralogs and three type 1 paracaspase paralogs in the vertebrate lineage both seem to have arisen in the last common ancestor of jawed vertebrates, which coincides with the evolution of the vertebrate adaptive immune system (Rast and Buckley 2013). Lampreys are in this respect more similar to invertebrates and only seem to have a single ancestral CARD-CC (Figure 4A) and a single type 1 paracaspase, a PCASP3 ortholog which is related to the parent of the PCASP3 and PCASP(1/2) branches in jawed vertebrates (Figure 1C, S1). Surprisingly, the supposedly ancestral CARD-CC in lampreys is clustering closer to CARD11 than CARD14 (Figure 4A). The invertebrate CARMA/CARD9-related CARD-CC domain proteins show a phylogenetic distribution similar to Bcl10 (Figure 4A), indicating that the entire CARD-CC/Bcl10/MALT1-like paracaspase (CBM) complex is co-evolving and that species with Bcl10-independent type 1 paracaspases rely on a completely independent activation mechanism. As seen in Bcl10, also the corresponding basic residues in CARD-CC required for Bcl10 interaction (Li et al. 2012) appear to be conserved as far back as *N. vectensis*. To functionally verify the conservation of an upstream CARD-CC interaction with the Bcl10/paracaspase complex, we co-expressed either human CARD9 or *Nematostella* CARD-CC together with human MALT1 in MALT1 deficient HEK293T cells to test their ability to induce MALT1-dependent NF-κB activation (Figure 4B). Also the cnidarian CARD-CC induced a MALT1-dependent NF-κB induction via endogenous human Bcl10. As expected from a conserved function of cnidarian Bcl10 (Figure 2B) and CARD-CC (Figure 4B) to activate MALT1-dependent NF-κB in human cells, cnidarian CARD-CC and Bcl10 were also found to enhance activated PKC signaling in human cells (Figure 4C). This indicates that the whole PKC-CBM signaling pathway is likely to have ancient evolutionary roots. Various PKC homologs can be found in a wide range of invertebrates (Kruse et al. 1996) and are not correlating with the presence of the CBM complex, reflecting their importance in many alternative pathways. Analogously, the CBM-interacting proteins AIP (Schimmack et al. 2014), Akt (Narayan et al. 2006), CaMKII (Ishiguro et al. 2007), caspase-8 (Kawadler et al. 2008), β-catenin and its destruction complex (Bognar et al. 2016), cIAP1/cIAP2 (Yang et al. 2016), CK1α (Bidère et al. 2009), CRADD (Lin et al. 2012), CSN5 (Welteke et al. 2009), MIB2 (Stempin et al. 2011), NOTCH1 (Shin et al. 2014), p62/SQSTM1 (Paul et al. 2014), RLTPR (Roncagalli et al. 2016) and Rubicon (Yang et al. 2012) also show a wide phylogenetic distribution with no indications of a CBM co-evolution. In contrast, A20 (Coornaert et al. 2008), DEPDC7 (D’Andrea et al. 2014), HECTD3 (Li, Chen, et al. 2013), LRRK1 (Morimoto et al. 2016) and Net1 (Vessichelli et al. 2012) show a phylogenetic distribution or BLASTp ranking that correlates with presence of the CBM complex. Other CBM interacting proteins like ADAP (Medeiros et al. 2007), BINCA (Woo et al. 2004), CKIP1 (Sakamoto et al. 2014), RIPK2 (Ruefli-Brasse et al. 2004) and USP2a (Li, He, et al. 2013) are poorly conserved in invertebrates and might represent more recently evolved CBM interaction partners. Taken together, we can, however, conclude that the core CBM complex components seem to be evolutionarily linked and functionally interacting ever since the last common ancestor of the planulozoans.

**Figure 4:**
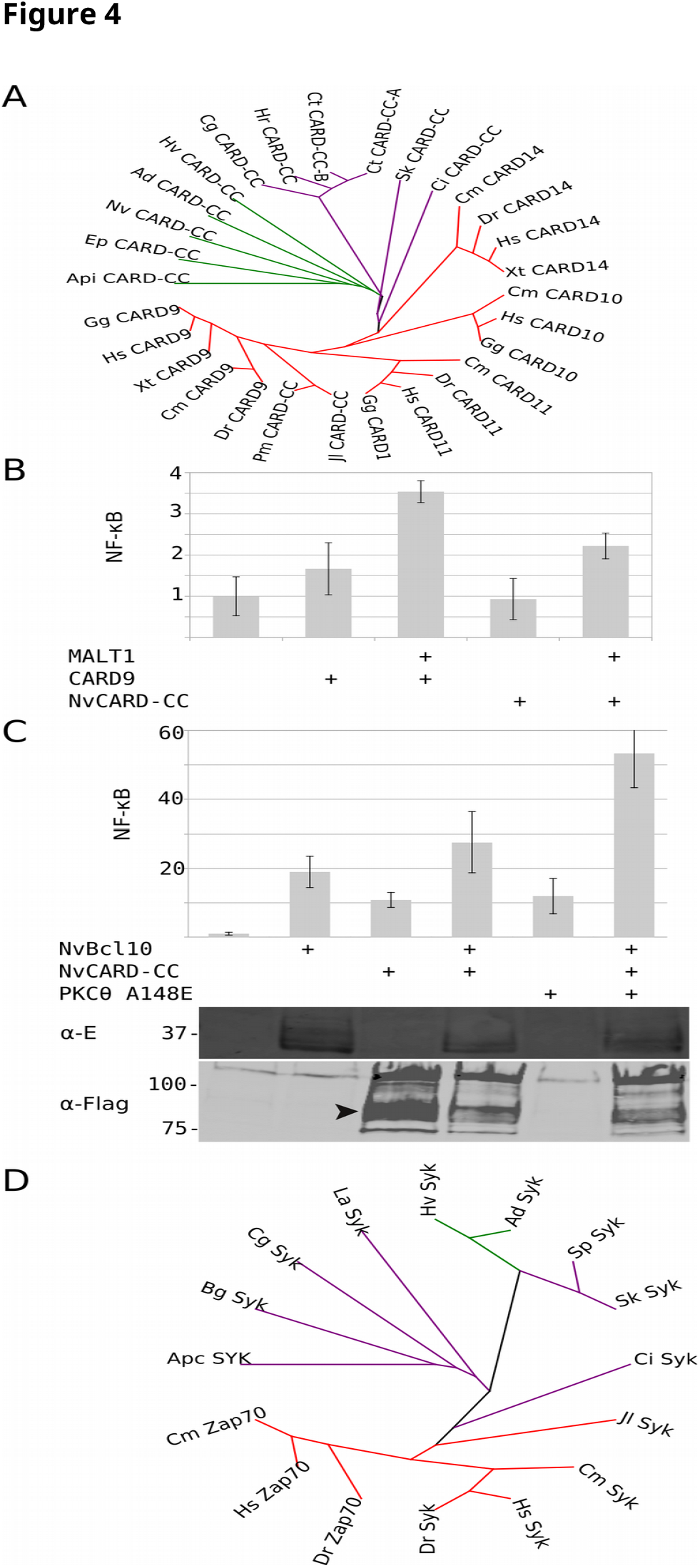
Evolution of CARD9/CARMA-homologs and upstream Syk. **A)** Maximum likelihood phylogeny (MUSCLE+PhyML) showing the relationships between CARD9, the three CARMA paralogs and their invertebrate CARD-CC homologs. **B)** Functional conservation of the cnidarian CARD-CC protein in MALT1-dependent NF-κB induction. Human CARD9 and *N. vectensis* CARD-CC were expressed in MALT1 deficient HEK293T cells together with an NF-κB-dependent luciferase reporter expression plasmid and a constituitively expressed β-galactosidase reporter gene. Where indicated human MALT1 was also co-expressed. Luciferase values are normalized against β-galactosidase and expressed as fold induction compared to samples not expressing CARD9 or CARD-CC (EV). Error bars represent 95% confidence intervals (Student’s t-distribution). CARD9 and *N. vectensis* CARD-CC induce NF-κB via MALT1, which is illustrated by the increase of luciferase activity when the cells are reconstituted with human MALT1. Experiments were repeated at least twice. **C)** Synergistic NF-κB induction by cnidarian CARD-CC and Bcl10 in presence of constituitively activated PKδ(A148E) in wild-type HEK293T cells. *N. vectensis* CARD-CC, Bcl10 and human PKδ(A148E) were expressed as indicated in HEK293T cells together with an NF-κB-dependent luciferase reporter expression plasmid and a constituitively expressed β-galactosidase reporter gene. Luciferase values are normalized against β-galactosidase and expressed as fold induction compared to samples not expressing CARD9 or CARD-CC (EV). Error bars represent 95% confidence intervals (Student’s t-distribution). The lower part of the panel shows the expression of *N. vectensis* Bcl10 as revealed by anti-E-tag development and *N. vectensis* CARD-CC as revealed by anti-Flag development on a western blot. Black arrowhead indicate expected protein size of *N. vectensis* CARD-CC (~86 kDa). Experiments were repeated at least twice. **D)** Maximum likelihood phylogeny (MUSCLE+PhyML) of Zap70/Syk homologs. Species key: <u>Vertebrates</u>: Hs=Human, Gg=Chicken, Xt=African clawed frog, Dr=Zebrafish, Cm=Elephant shark, Pm=Sea lamprey, Jl=Japanese lamprey <u>Tunicates</u>: Ci=vase tunicate <u>Hemichordates</u>: Sk=Acorn worm <u>mollusks</u>: Cg=Pacific oyster <u>Annelids</u>: Ct=polychaete worm, Hr=leech <u>Cnidaria</u>: Nv=Starlet sea anemone, Hm=Hydra, Ep=Sea anemone. Red branches highlight vertebrate sequences, green branches cnidaria and purple branches bilaterian invertebrates with Bcl10.

### ITAM receptors and Syk : a potential original pathway for the CBM complex

Given the CARD9-like domain organization of the invertebrate and lamprey CARD-CC homologs, it is tempting to speculate that the ancestral and invertebrate CARD-CC proteins play a similar role. CARD9 interacts with the intracellular pathogen-associated molecular pattern receptor Nod2 and is critical for downstream MAP kinase responses (Parkhouse et al. 2014). Nod2 originated in vertebrates (Boyle et al. 2013), but is a member of the NLR receptor family where metazoan homologs can be found at least as far back as sponges (Yuen et al. 2014). The metazoan NLR family is clearly ancient and has a domain structure which is intriguingly similar to fungal sexual compatibility proteins and a large family of plant pathogen resistance (R) proteins, but this similarity is most likely due to convergent evolution (Staal and Dixelius 2007). At this moment, no role for Bcl10 or MALT1 has been found in Nod2 signaling, indicating that this is a CARD9-specific function. CARD9 and Bcl10 are also critical components in RIG-I mediated antiviral signaling (Poeck et al. 2010). RIG-I is also highly conserved and can be found as far back as cnidarians and with top scoring BLASTp hits in invertebrate species that have CARD-CC and Bcl10 (full-length hits in deuterostomes and lophotrochozoans), which makes it an interesting candidate original receptor pathway. RIG-I has also been shown to be functionally conserved as far back as mollusks (Zhang et al. 2014). The most prominent role of CARD9 is however in a CBM complex in the evolutionary conserved C-type lectin signaling pathway (Sattler et al. 2012; Drummond and Lionakis 2016). CARD9 is critical for C-type lectin-mediated immunity against fungal infections in humans, which is also the most clinically relevant effect of CARD9 deficiency (Alves de Medeiros et al. 2016). Interestingly, a survey of invertebrate Dectin-1 C-lectin domain homologs finds back top-scoring hits from mollusks and cnidarians but not the much better characterized arthropod and nematode genomes (Pees et al. 2016), similar to previous observations (Wood-Charlson and Weis 2009). Recently, the vav1-3 proteins were found to be critical for CARD9 signaling downstream of lectin receptors (Roth et al. 2016), and also these proteins are more similar in cnidarians and lophotrochozoans compared to other invertebrates. C-type lectins are already associated to innate immunity in mollusks (Li et al. 2015) and cnidarians (Vidal-Dupiol et al. 2009). Human Dectin-1 signals to the CARD9/Bcl10/MALT1 complex via the tyrosine kinase Syk and PKCδ. The most similar invertebrate Syk/Zap70 homologs also correlate with the presence of Bcl10/CARD-CC, where especially the N-terminal sequence of Syk was specific for those organisms (Figure 4D). This is in agreement with earlier observations of a loss of the Syk kinase during metazoan evolution (Steele et al. 1999). Strikingly, the pattern of Syk-containing organisms is largely overlapping with organisms containing Bcl10 and CARD-CC homologs, with a duplication event of Syk to Zap70 in the jawed vertebrates. The 200 first residues of Syk or Zap70 made the CARD-CC/Bcl10-correlated phylogenetic distribution even clearer with deuterostome, mollusk and cnidarian proteins among the top invertebrate hits. The N-terminal SH2 domains in Syk and Zap70 are critical for interaction with upstream ITAM domain containing receptors (Flaswinkel et al. 1995; Mócsai et al. 2010). The CBM-linked phylogenetic distribution of the SH2 domains in the tyrosine kinase Syk indicates ITAM containing upstream receptors linked to the CBM complex. In contrast, another invertebrate SH2 domain tyrosine kinase ("Shark"), which has been shown to also mediate ITAM-dependent immune-related signals (Ziegenfuss et al. 2008), does not show a correlation with the CBM complex components. Furthermore, there is no sequence hit of the N-terminal ITAM - containing intracellular domain of Dectin-1 in mollusks or cnidarian transmembrane C-type lectins and there is currently no proof of a C-lectin receptor/Syk pathway in invertebrates. Hypothetically, invertebrate C-type lectin receptor signaling could, however, utilize an ITAM/Syk dependent pathway via β-integrin (Jakus et al. 2007; Wang et al. 2014), but the vertebrate ITAM adaptors Dap12 and FcRγ for integrin signaling are not conserved. The ITAM - dependent signaling in invertebrates could also be mediated by another class of receptors, as suggested for lamprey (Liu et al. 2015). If the RIG-I/CBM or ITAM/Syk/PKC/CBM pathways are shown to be conserved, further insight on the biology and regulation of the CBM complex would not only benefit biomedical research against (auto)immune diseases and cancer (Demeyer et al. 2016), but could also impact a wide range of other areas such as mollusk (aqua)culture and environmentally important challenges like the host immunity component of coral bleaching (Vidal-Dupiol et al. 2009; Bosch et al. 2014). Interestingly, a MALT1 homolog was found to be one of the most significantly up-regulated genes in corals after natural bleaching stress (Pinzón et al. 2015). For now we can only conclude that the intriguing patterns of co-evolution can correspond to an evolutionary conserved functional interaction. These observations can serve as foundations for testable hypotheses in future functional characterizations of the CBM complex and its associated proteins and pathways in alternative species.

### Overlapping expression domains of CBM complex components in the starlet sea anemone (Nematostella vectensis)

Since the *N. vectensis* type 1 paracaspase paralog “B” was found to interact with *N. vectensis* and human Bcl10 (Figure 2D) and *N. vectensis* CARD-CC showed synergistic activity with *N. vectensis* Bcl10 (Figure 4C), it is likely that these three components form a CBM signaling complex. To form a complex, the different genes need to be co-expressed. Investigations of the expression patterns of the two type 1 paracaspase paralogs, Bcl10 and CARD-CC during *N. vectensis* embryo development (11 stages, from unfertilized egg to juvenile stage, 14 days post fertilization) using *in situ* hybridization (Genikhovich and Technau 2009) revealed overlapping temporal and spatial expression domains of the proposed CBM complex components (Figure 5). The assumed temporal overlap is based on a visual interpretation from a series of developmental snap shots (Gombrich 1980), but the developmental variation present in each stage sample helps the interpretation. Stainings revealed similar expression patterns for all four genes. All four genes showed expression from late planula and onwards (Figure 5) roughly corresponding to *Nvncam2* (Marlow et al. 2009), *Nvmef2-II* (Genikhovich and Technau 2011) and *Nvdelta* (Marlow et al. 2012), indicating neuronal expression. *NvBcl10, NvPcasp-t1B* and *NvCard-cc* also show an expression pattern similar to *Nvmef2-II* in gastrula. In addition, all four genes also show high expression in the tentacles in the later stages. The expression pattern of *N. vectensis* Bcl10 is intriguing and might indicate an ancient conserved developmental role, since it has been shown that Bcl10 is involved in neural tube closure in mouse (Ruland et al. 2001). Strong neuronal expression profiles of MALT1 (EMAGE: 12681) during mouse development (Richardson et al. 2013) also indicate that a complete CBM complex might be involved in this process. On the other hand, no neurodevelopment defects were found in MALT1 deficient mice (Ruland et al. 2003). Cnidaria do not have a central nervous system and no obvious homolog of the neural tube in chordates or the corresponding structures in other deuterostomes (Nielsen 2015). However, if a conserved role in cnidarian neuronal development can be established, this would indicate that Bcl10 and possibly the other CBM complex components would have evolved together with the evolution of the nervous system (Kelava et al. 2015), and this could represent one of the original functions of the CBM complex. In case there is a functional involvement of CBM complex components in the cnidarian nervous system, the reported interaction of the CBM complex with Notch1 would be a likely conserved mechanism (Marlow et al. 2012; Shin et al. 2014). The *N. vectensis* type 1 paracaspase paralog “A”, which failed to show any Bcl10 interaction (Figure 2D), also showed a similar expression pattern, indicating that the Bcl10-independent type 1 paracaspases found in nematodes and arthropods might have a similar function.

**Figure 5:**
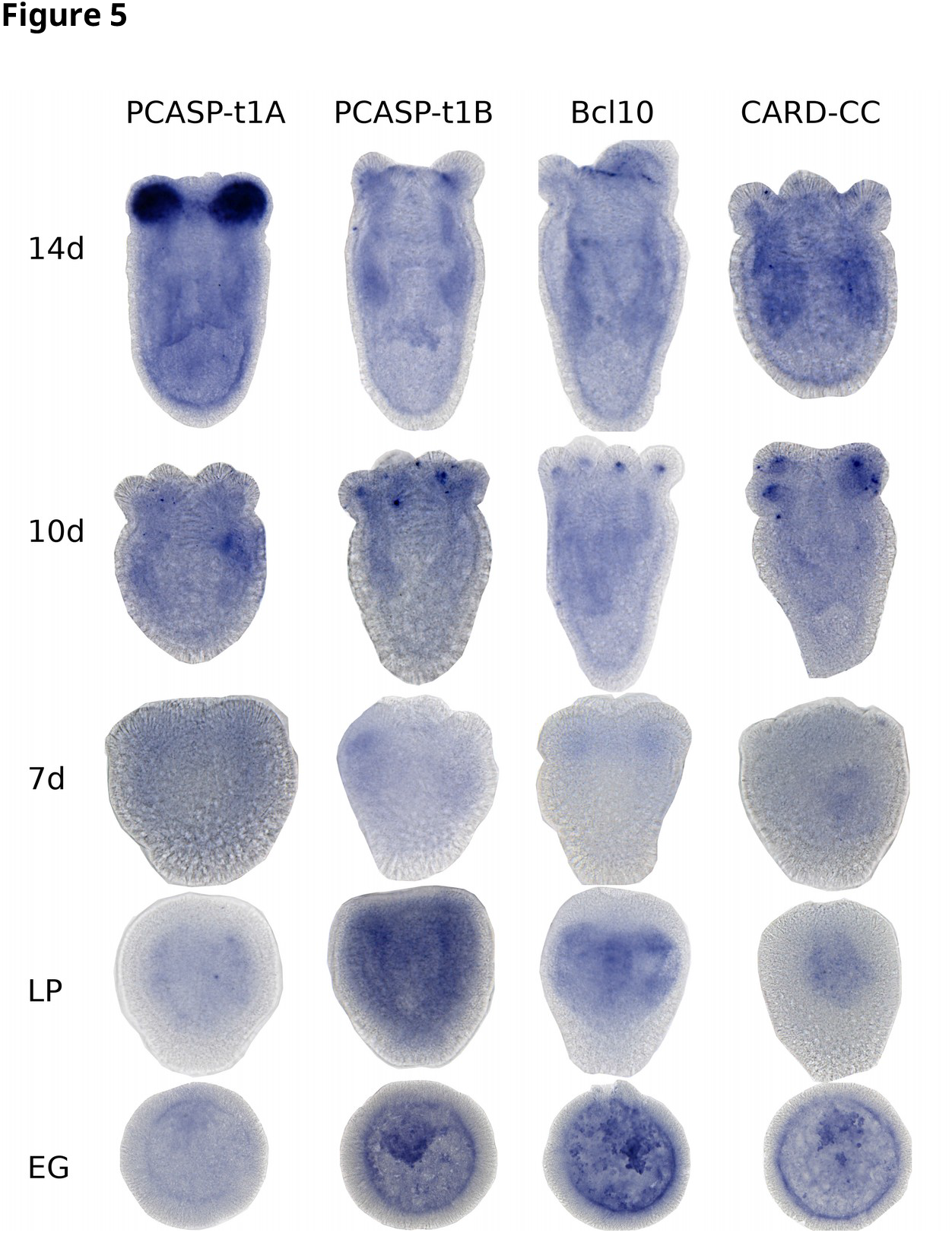
The CBM complex components show overlapping expression domains in N.vectensis. Developing *N. vectensis* embryos were stained for the indicated genes with in situ hybridization at 11 different developmental stages, from unfertilized egg to 14 days post fertilization. Informative pictures from 14 days (14d), 10 days (10d), 7 days (7d) late planula (LP) and early gastrula (EG) for all 4 CBM complex genes are shown.

### CBM and NF-κB independent functions of type 1 paracaspases

The nematode model organism *C. elegans* is a promising system to specifically investigate unconventional functions of type 1 paracaspases because it lacks CARD-CC, Bcl10 and NF-κB. Despite the lack of known upstream and downstream proteins in the signaling pathway, the WormBase *C. elegans* phenotype database indicates an important role for the type 1 paracaspase with a “lethal or sterile” mutant phenotype (*tm289* vs *tm321*) in nematodes (C. elegans Deletion Mutant Consortium 2012). To investigate this further, we analyzed lifespan of *C. elegans* in which the paracaspase *F22D3.6* was knocked down by RNAi. We silenced this gene systemically in the wild type strain N2 (all tissues except the neurons) and in a strain hypersensitive to neuronal RNAi (TU3311). While systemic knockdown of *F22D3.6* showed no effect on lifespan, strong silencing of this gene in the neurons caused a slight but significant reduction of lifespan (P< 0.05 in all 3 replicates, Figure 6 A, B), hinting at a vital role of this paracaspase homolog in the *C. elegans* neurons. Remarkably, paracaspase knockdown in the neuronal RNAi hypersensitive strain caused increased motility (Figure 6C). This highlights an important role for CBM-independent type 1 paracaspase functions and indicates that further investigations on the role and function of *F22D3.6* in *C. elegans* could be highly interesting. Especially phenotypic characterization of a paracaspase protease-dead knock-in mutant and identification of *C. elegans* paracaspase interacting proteins could help defining its functions in this organism. Ultimately, identifying the CBM-independent type 1 paracaspase activation mechanism in organisms like *C. elegans* could potentially also lead to the discovery of a novel fifth (Bcl10-independent) MALT1 activation mechanism, apart from the four (CARD9, 10, 11, 14) currently known CBM complexes in humans.

**Figure 6:**
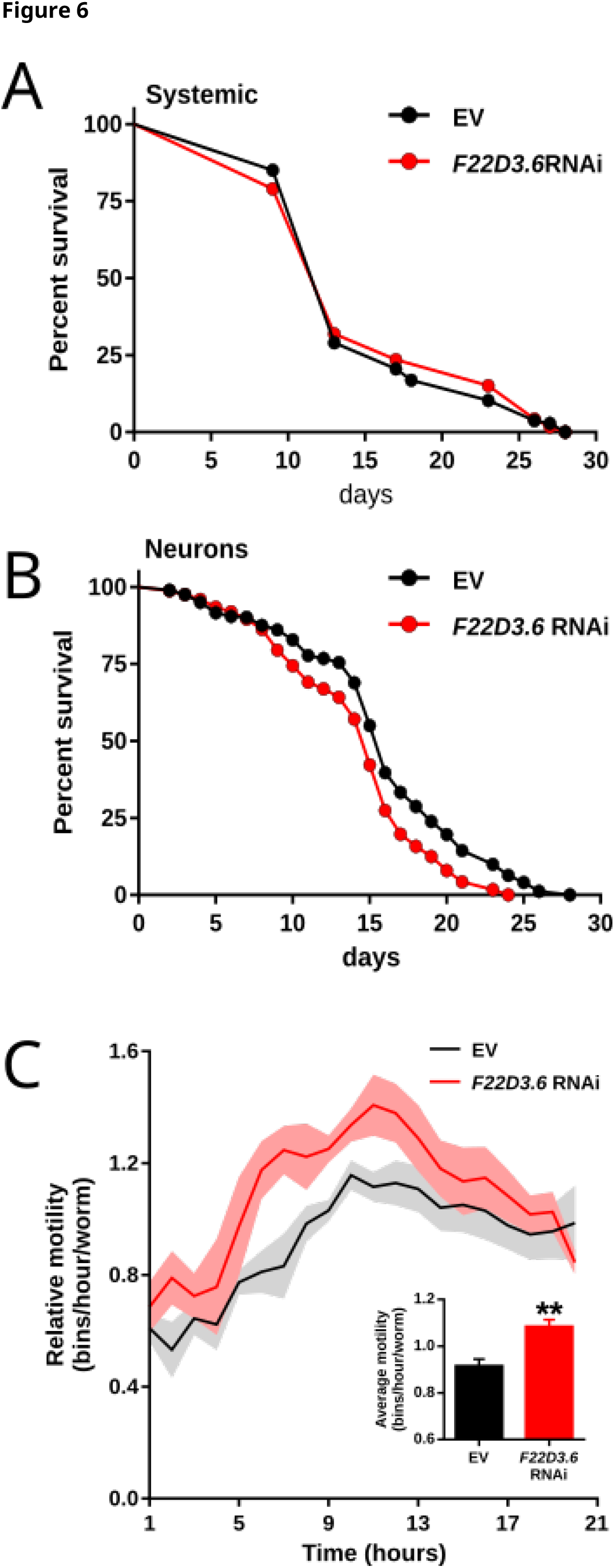
CBM and NF-κB independent functions of the type 1 paracaspase F22D3.6 in *C. elegans*. **A)** Wild-type (N2 strain) *C. elegans* worms were fed *E. coli* expressing RNAi targetting *F22D3.6* and the worms were regularly monitored for viability compared to worms fed control RNAi. RNAi is typically targeting every cell except neurons in wild-type worms. **B)** Neuronal RNAi importer transgenic (TU3311 strain; *unc-119p::sid-1* transgenic) *C. elegans* worms were fed *E. coli* expressing RNAi targetting *F22D3.6* and the worms were regularly monitored for viability compared to worms fed control RNAi. **C)** Neuronal RNAi importer transgenic (TU3311 strain; *unc-119p::sid-1* transgenic) *C. elegans* worms were fed *E. coli* expressing RNAi targetting *F22D3.6* and the worms were monitored for motility compared to worms fed control RNAi using the Wmicrotracker device. The figures A-C represent pooled results of three biological replicates carried out in 20°C.

### Future challenges

With the observation that mollusk paracaspases have conserved protease activity and specificity, but fail to induce NF-κB in a human cellular background, we are starting to unravel the sequence of evolutionary events leading to the current MALT1 protease- and NF-κB-activating scaffold activities in humans. Our results indicate that the MALT1-like protease activity and substrate specificity precedes the evolution of MALT1-like NF-κB-activating scaffold function. The evolutionary origin of the scaffold function appears to be elusive, with a lack of conservation of TRAF6 binding motifs among vertebrate paracaspases that are able to induce NF-κB. We still don’t know how far back MALT1-like activities such as TRAF6 interaction and NF-κB induction, protease activity and specificity are conserved. It would thus be very interesting to investigate the functional properties of paracaspases from species closely related to the jawed vertebrates (e.g. lampreys, tunicates, lancelets and hemichordates) to identify when type 1 paracaspases evolved into NF-κB-inducing scaffold proteins. A major future challenge will be to collect and functionally evaluate more invertebrate type 1 paracaspase, Bcl10 and CARD-CC homologs to verify the proposed correlation of a CARD-CC/Bcl10-dependent ancestral type 1 paracaspase paralog with MALT1-like activity and to model the evolution of the MALT1/Bcl10/CARD-CC interaction. There are already very good structure-function studies done on the human Bcl10/MALT1 interaction (Langel et al. 2008) and the CARMA1/Bcl10 interaction (Li et al. 2012). Functional characterization of evolutionary distant homologs do however provide complementary insights (Hopf et al. 2014). While traditional structure-function analyses mutate single residues predicted to be important in order to disrupt activity, our identification of functionally conserved distant homologs meant that we could simultaneously “mutate” up to 65% of all residues in the critical interaction domain without disrupting activity. Patterns of co-evolution of interacting proteins does not only give information on single protein-protein interactions, since it also can provide information on a higher abstraction level for entire signaling networks (Cui et al. 2009). There are however several aspects of CBM evolution that are not yet clear. For instance, no Bcl10 or CARD-CC homolog is currently found in lancelets, which clearly have a *Pcasp3* ortholog that is supported by synteny (Hulpiau et al. 2016). The limited number of invertebrate true *Bcl10* homologs that can be identified in public sequence data is currently a limitation for further analysis. CRADD homologs are often picked up as false positives in distant species since they contain a CARD domain that is very similar to Bcl10 (Lin et al. 2012; Qiao et al. 2014). The current model proposes an ancient parallel evolution of a Bcl10-dependent and a Bcl10-independent paracaspase (Figure 7A). An alternative scenario is that Bcl10-independence has evolved several times. To further clarify this, more invertebrate sequences from informative phyla are needed (GIGA community of scientists 2014). Several proteins associated to MALT1 in humans are conserved as far back as cnidarians, such as CARMA/CARD9 (CARD-CC), Bcl10, TRAF6, TRAF2 and CYLD (Hulpiau et al. 2016), and we have now shown that many are functionally conserved in a human cellular environment. Investigating early-diverging biological model systems such as the cnidarians for protein interactions and signal transduction mechanisms could further highlight the original and most conserved functions in a native context. Recent advances in cnidarian cell culture might enable such functional analysis by transient overexpression (Rabinowitz et al. 2016). Also CRISPR/Cas9 is now a viable option in many invertebrate model systems for generation of knock-out or knock-in mutant animals (Ikmi et al. 2014; Chen et al. 2016). The sea anemone cnidarian model organisms *N. vectensis* and *Aiptasia* might, however, not be the best choices because they express a short form of CYLD and do not have one of the typical cnidarian type 1 paracaspase paralogs found in hydra and corals (PCASP-t1C, Figure 1C, S1). It is possible that the type 1 paracaspases from hydra or coral have more MALT1-like characteristics and cleave the full-length cnidarian CYLD. Furthermore, corals (e.g. *Acropora digitifera*) have been found to have a massively expanded immune repertoire compared to *N. vectensis* (Shinzato et al. 2011; Quistad et al. 2014). The paracaspase expression patterns in *N. vectensis* and mice and the phenotypic effects of paracaspase silencing in *C. elegans* indicate a neuronal role for type 1 paracaspases. Because of the possible neuronal connection, also the mollusk *Aplysia* neuronal model systems could be interesting for functional investigations of the CBM complex components.

**Figure 7:**
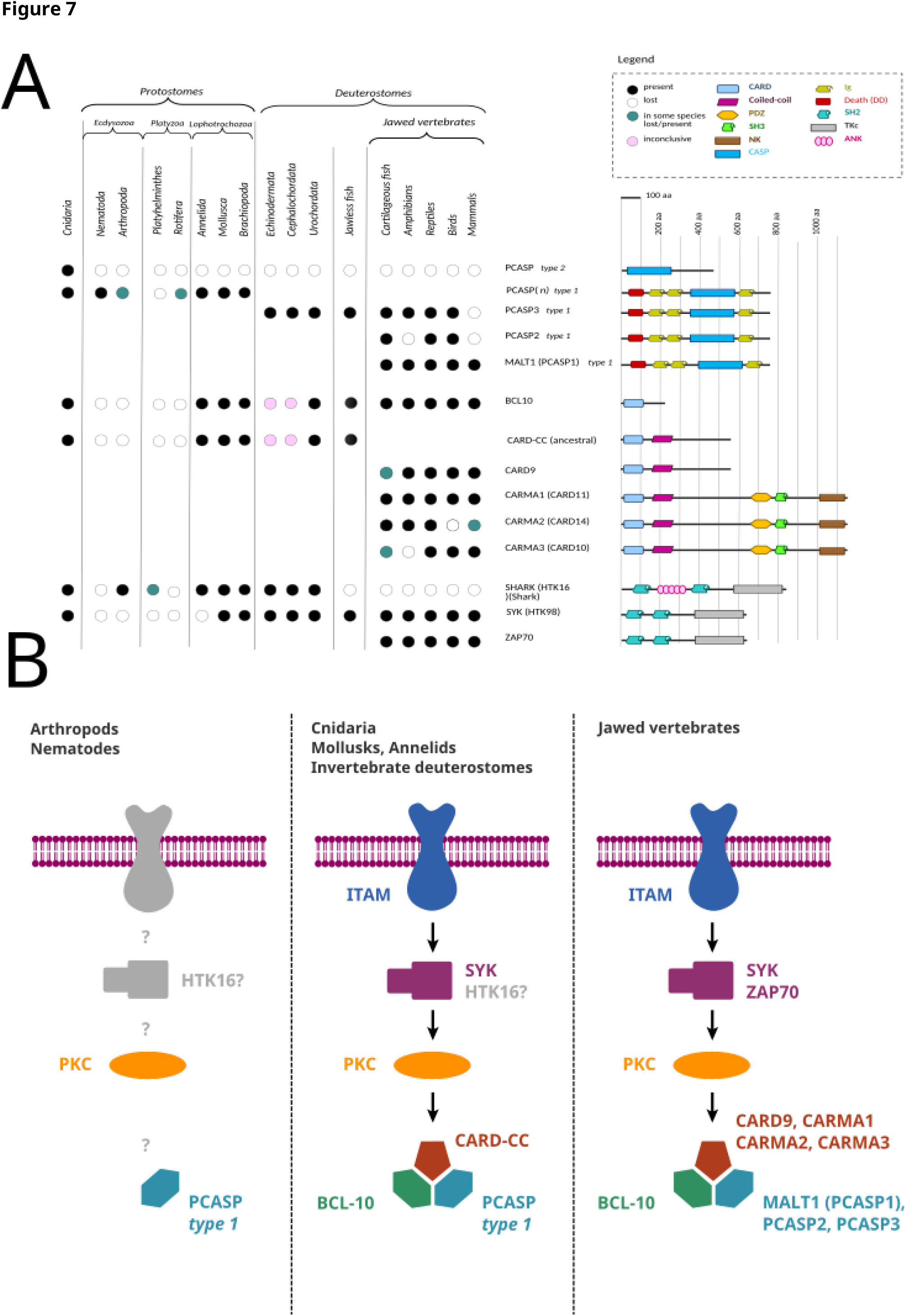
Co evolution and proposed signaling model. **A)** Patterns of co-evolution of Syk and the CBM complex components in various organisms. Type 1 paracaspases prior to Deuterostomes are annotated as PCASP(n) since currently available sequences cannot determine whether a distant invertebrate paracaspase is an ancient PCASP3 paralog or ortholog. One model proposes 2 ancient type 1 paracaspases, one Bcl10-dependent and one Bcl10-independent. The CARD-CC/Bcl10-dependent type 1 paracaspase shows MALT1-like activities. Deuterostomia (including tunicates, lancelets, vertebrates and hemichordates), annelids and mollusks inherited the Bcl10-dependent type 1 paracaspase whereas most other bilaterian invertebrates kept the Bcl10-independent type 1 paracaspase. The model is based on currently available reliable sequence information and might change with additional data. Analogously, CARD-CC became duplicated and later fused with MAGUK domains in the jawed vertebrates. At this moment, we uncertain which of the 4 jawed vertebrate CARD-CC paralogs (CARD9, 10, 11, 14) should be considered ortholog of the ancestral CARD-CC. Also upstream Syk became duplicated in jawed vertebrates, resulting in Zap70. **B)** Proposed signaling model in various organism classes. Nothing is known about upstream activators of type 1 paracaspases in CARD-CC / Bcl10-independent organisms such as arthropods and nematodes.

## Materials & Methods

### Sequences of type 1 paracaspases, Bcl10 and CARD-CC homologs

Protein sequences of type 1 paracaspase, Bcl10 and CARMA/CARD9 homologs were retrieved from NCBI (https://www.ncbi.nlm.nih.gov), Ensembl (http://metazoa.ensembl.org), JGI
(http://genome.jgi.doe.gov/), OIST marine genomics (http://marinegenomics.oist.jp) (Shinzato et al. 2011) (Luo et al. 2015) (Simakov et al. 2015), ReefGenomics (http://reefgenomics.org/)(Baumgarten et al. 2015) and ICMB (https://imcb.a-star.edu.sg) (Mehta et al. 2013; Venkatesh et al. 2014) using BLASTP (Johnson et al. 2008). All sequences used in the analyses can be found in supplemental material for independent replication of the results (supplemental texts 1–4).

### Sequence alignment and phylogenetic analysis

Sequence alignment was performed on the full sequence, using the different alignment algorithms Clustal Omega (http://www.clustal.org/omega/) (Sievers and Higgins 2014), MUSCLE (http://www.drive5.com/muscle/) (Edgar 2004), and T-coffee (http://www.tcoffee.org/) (Notredame et al. 2000). Phylogenetic analysis was performed with PhyML (http://atgc_montpellier.fr/phyml/) (Guindon et al. 2009) and MrBayes (http://mrbayes.sourceforge.net/) (Ronquist and Huelsenbeck 2003) methods. N-terminal paracaspase sequences were trimmed out from a MUSCLE multiple sequence alignment using Jalview (Waterhouse et al. 2009). Trimmed N-terminal paracaspase sequences less than 190 residues long were excluded from the phylogenetic analysis. Both alignments and phylogenetic analyses were performed using UGENE (http://ugene.net/) (Okonechnikov et al. 2012) on Arch (http://www.archlinux.org) Linux (Torvalds 1999). For the figures, one of the most representative trees (alignment and phylogenetic analysis) were selected. For independent replication of the results, all sequences used in the phylogenetic analysis are available in the supplemental data (supplemental texts 1–4). The final trees were re-drawn to unrooted radial cladograms using dendroscope (https://dendroscope.org) (Huson and Scornavacca 2012). Metadata by coloring the branches was manually added using Inkscape (https://inkscape.org).

### RNA isolation and cDNA synthesis

RNA of *N. vectensis* from various developmental stages was isolated with TRIzol (Thermo Fisher) and pooled. 1.5 μg of total RNA was subjected to 5’-RACE with a GeneRacer kit (Invitrogen) according to the manufacturer’s protocol. Briefly, RNA was treated with calf intestinal phosphatase (CIP) to remove the 5’ phosphates from truncated RNAs and non-mRNAs. After dephosphorylation of the RNA with tobacco acid pyrophosphatase (TAP), lyophilized GeneRacer RNA Oligo (provided in the kit) was then added to the 5’ end of the RNA with RNA ligase. The ligated RNA was reverse transcribed to cDNA using superscript III with random primers and used as templates for PCR amplifications.

### Cloning of invertebrate homologs

Plasmids of the cloned genes were deposited in the BCCM/LMBP plasmid collection along with detailed descriptions of cloning strategy and plasmid sequence (http://bccm.belspo.be/about-us/bccm-lmbp). The starlet sea anemone (*N. vectensis*) type 1 paracaspase paralog “A” (LMBP: 9589) and zebrafish PCASP3 (LMBP: 9573) were cloned previously (Hulpiau et al. 2016). The *N. vectensis* type 1 paracaspase paralogs “A” (LMBP: 9636) and “B” (LMBP: 9825) and pacific oyster (*Crassostrea gigas*, LMBP: 9826) were cloned behind the human ETV6 HLH domain for dimerization-induced activation as described previously (Malinverni et al. 2010; Baens et al. 2014; Hulpiau et al. 2016). Human (LMBP: 9637), zebrafish (LMBP: 9665), pacific oyster (LMBP: 9666) and *N. vectensis* (LMBP: 9822) Bcl10 were cloned in the pCAGGS vector with an N-terminal E-tag. For functional characterization of the Bcl10 CARD domain, hybrid Bcl10 clones where human residues 1-102 were replaced with the corresponding sequence from pacific oyster (CgN, LMBP: 10190) or *N. vectensis* (NvN, LMBP: 10191) were cloned. Also minimal active Δ119-223 clones of human Bcl10 (LMBP: ), pacific oyster (CgN, LMBP: ) and *N. vectensis* (NvN, LMBP: ) hybrid Bcl10 were used for functional tests. *N. vectensis* CYLD (LMBP : 9900) was also cloned in pCAGGS with an N-terminal E-tag. The *N. vectensis* homologs of CARD-CC (LMBP: 9854) TRAF6 (LMBP: 9855) and TRAF2 (LMBP: 9856) in a pCDNA3 vector with N-terminal Flag tag. For further specific cloning purposes, human CARD9 (LMBP: 9877), Bcl10 (LMBP: 9872), MALT1 (LMBP : 9104, 9105) and *N. vectensis* CARD-CC (LMBP: 9873), Bcl10 (LMBP: 9874), PCASP-t1A (LMBP : 9875) and PCASP-t1B (LMBP : 9876) were cloned into gateway-compatible pENTR-vectors.

### Cell culture, transfection and expression analysis

MALT1 deficient HEK293T cells (clone #36) (Hulpiau et al. 2016) were grown under standard conditions (DMEM, 10% FCS, 5% CO_2_, 37 °C) and transfected with the calcium phosphate method (Anon 2005). For evaluation of the conservation of cleavage activity, the HLH-fused paracaspase constructs were co-transfected with wild-type CYLD (LMBP: 6613) or the uncleavable CYLD-R324A (LMBP: 6645) mutant. Cells transfected for cleavage activity evaluations were lysed directly in Laemmli buffer (0.1% 2-Mercaptoethanol, 5ppm Bromophenol blue, 10% Glycerol, 2% SDS, 63 mM Tris-HCl (pH 6.8)). For evalutation of conservation of NF-κB induction, the HLH paracaspase fusions were co-transfected with a NF-κB-dependent luciferase reporter expression plasmid (LMBP: 3249) and an actin promoter-driven β-galactosidase expression plasmid (LMBP: 4341) as transfection control. The cells used for luciferase analysis were washed with PBS and lysed in luciferase lysis buffer (25mM Tris pH7.8, 2mM DTT, 2mM CDTA, 10% glycerol, 1% Triton X-100). For the colorimetric determination (at 595nm) of β-galactosidase activity, chlorophenol red-β-D-galactopyranoside (CPRG) (Roche diagnostics) was used as a substrate. Luciferase activity was measured by using beetle luciferin (Promega) as a substrate and luminescence was measured with the GloMax^®^ 96 Microplate Luminometer (Promega). Luciferase data processing and calculation of 95% confidence intervals (Student’s t-distribution (Student 1908)) was done in LibreOffice (www.libreoffice.org) Calc (Gamalielsson and Lundell 2014). For evaluation of the functional conservation of Bcl10 homologs, the Bcl10 clones were co-transfected with the NF-κB luciferase reporter and β-galactosidase expression plasmids in the MALT1 deficient HEK293T cells with or without reconstitution with human MALT1 (LMBP: 5536). Human CARD9 (LMBP: 9609) was used as control for evaluations of the functional conservation of CARD-CC proteins. For evaluation of PKC activation of *Nematostella* CARD-CC/Bcl10 interactions, the constituitively activated human PKCθ(A148E) (LMBP: 8925, 9045) was used. Detection of cleaved CYLD was done with the E10 antibody (Santa Cruz Biotechnology) recognizing the C-terminal 70kDa cleavage band or anti-E-tag (ab66152, Abcam) recognizing the 40kDa N-terminal cleavage band. Expression of the fused paracaspases was determined with anti-Flag (F-3165, Sigma). Human MALT1 was detected by the EP603Y monoclonal rat antibody (Abcam) and the E-tagged Bcl10 clones with anti-E-tag. All western blots were developed on an Odyssey scanner (LI-COR).

### Yeast-2-hybrid assay

Human MALT1 (LMBP: 9880, 9899), Bcl10 (LMBP : 9879, 9885), CARD9 (LMBP: 9878, 9884) and Nematostella PCASP-t1A (LMBP: 9898) PCASP-t1B (LMBP: 9883, 9888), Bcl10 (LMBP : 9882, 9887) and CARD-CC (LMBP: 9881, 9886) were cloned into the pdGADT7 or pdGBKT7 vectors by Gateway LR reaction. The ENTR vectors were linearized to enable cloning into the kanamycin-resistant destination vector without background contamination. The Matchmaker Gold Yeast Two-Hybrid System (Clontech) was used with the Y2H Gold yeast strain to investigate protein-protein interactions. A pre-culture was made the day before transformation, by inoculating about 10 colonies of Y2H gold strain in 5 ml YPDA medium and growing it for about 4 h in a 30°C shaking incubator. The pre-culture was transferred to 35 ml YPDA and grown overnight in a 30°C shaking incubator. On the day of transformation, the overnight culture was diluted to an OD_600_ of 0.2 in YPDA (depending on the number of transformations, 10 ml YPDA/transformation) and grown in a 30°C shaking incubator until an OD_600_ of 0.6–0.8. After a 5 min centrifugation step at 2100 rpm 23 °C, the yeast pellet was resuspended in 10 ml Milli-Q water and centrifuged again for 5 min. After resuspending the pellet in 1× TE/LiAc, 100 μl of competent cells were mixed with 100 μg denatured salmon sperm DNA, 1 μg bait plasmid, 1 μg prey plasmid and 600 μl fresh PEG400/LiAc. The yeast-DNA mixtures were incubated in a 30°C shaking incubator for 30 min. The yeast cells were transformed via heat-shock at 42°C for 15 min. After a 1-min incubation on ice and a 30-sec centrifugation step, the pellet was resuspended in 1× TE and plated on minimal synthetic drop-out medium (SD) lacking leucine and tryptophan (SD -Leu/-Trp). After 4 days of incubation at 30°C, colonies were picked and incubated overnight in 200 μl SD/-Leu/-Trp medium in a 96-well plate. Transformed yeast cells were grown overnight in a 30°C incubator. Cultures were then stamped on SD/-Leu/-Trp and SD/-Leu/-Trp/-His/-Ade/+X-a-gal (40 μg/mL 5-bromo-4-chloro-3 indolyl-b-D-galactopyranoside) plates using an iron 96-well stamp and incubated for 3-7 days at 30°C until blue colonies were visible.

### In situ expression analysis in N. vectensis

As RNA probe templates, pDEST12.2 clones of *N. vectensis* CARD-CC (LMBP: 9908), Bcl10 (LMBP: 9902), PCASP-t1A (LMBP: 9903) and PCASP-t1B (LMBP: 9904) were generated by Gateway LR reaction. SP6 RNA polymerase (Promega) was used to generate labeled RNA probes. Fixed *N. vectensis* embryos were transferred into wells and rehydrated with 60% methanol / 40% PBS with 0.1% Tween 20 (PBSTw), 40% methanol / 60% PBSTw and four times with 100% PBSTw. The samples were then digested with 10 μg/ml Proteinase K (prepared in PBSTw) for 20 min. The reaction was stopped by two washes with 4 mg/ml glycine. The embryos were washed first with 1% triethanolamine (v/v in PBSTw), twice with 1% triethanolamine / 3 μl acetic anhydride and then twice with 1% triethanolamine / 6 μl acetic anhydride. After two washes with PBSTw, the embryos were refixed in 3.7% paraformaldehyde (v/v in PBSTw) for one hour and washed five times with PBSTw. Samples were prehybridized in 50% PBSTw / 50% Hybridization buffer (Hybe) (50% formamide, 5X SSC, 50 μg/ml heparin, 0.1% Tween 20 (v/v), 1% SDS (v/v) 100 μg/ml SS DNA and DEPC water) for 10 min, 100% Hybe buffer for 10 min and 100% Hybe buffer overnight at 60°C. Labelled RNA probes were diluted to 0.5 ng/μl Hybe buffer and denatured at 85°C for 10 min. Hybe buffer was removed from the embryos and for each reaction 250-300 μl working stock probe was added into the *in situ* plate. The sieves with embryos were transferred to the *in situ* plate and sealed to prevent evaporation. The embryos were then incubated at 60°C for 48 to 72 hours. The sieves were transferred to a clean rack filled with fresh preheated Hybe buffer and incubated at 60°C for 10 min. Then the samples were washed with 100% Hybe buffer and incubated at the hybridization temperature for 40 min. The embryos were washed at hybridization temperature for 30 min; once with 75% Hybe / 25% 2X SSCT (pH 7.0, 0.3 M sodium citrate, 3 M NaCl and 0.1% (v/v) Tween 20), once with 50% Hybe / 50% 2X SSCT, once with 25% Hybe / 75% 2X SSCT, once with 2X SSCT and finally three times with 0.05X SSCT. Prior to the blocking step, the samples were washed three times with 100% PBSTw (each 10 min) at room temperature. To decrease the unspecific background, the samples were blocked in Roche blocking reagent (supplemented with 1% (w/v) 1X maleic acid) for one hour at room temperature. The embryos were then incubated with antibody solution (Roche anti-digoxigenin-AP (alkaline phosphatase) diluted 1/2000 in blocking buffer) at 4°C overnight. The sieves were rinsed with blocking buffer and washed 10 times with 100% PBSTw (each 15 min). The embryos were developed in AP substrate solution (5 M NaCl, 1 M MgCl_2_, 1 M Tris pH 9.5 and 0.1% (v/v) Tween 20) at room temperature. Color development was checked every 10 min for 2 hours and AP substrate solution was replaced if an extended developing period was required. Once the probe development reached the desired level, the reaction was stopped by washing with 100% PBSTw. Next, the samples were washed with 100% ethanol for 1 hour and rinsed several times with 100% PBSTw. Finally, the specimens were washed with 85% glycerol (in PBSTw) at 4°C overnight and embedded to microscope slides using polyvinyl alcohol hardening mounting medium (10981-100ML, Sigma-Aldrich).

### Microscopy

Images were captured with a Axio Scan.Z1 (Zeiss). Images were acquired with a 20X Plan-Apochromat 0.8 NA dry objective, using a Hitachi HV-F202SCL camera.

### *RNAi silencing of F22D3.6 in* C. elegans *and phenotypic analysis*

SID-1 is a transmembrane protein, responsible for the passive uptake of dsRNA but this protein is only present in all cells outside the nervous system. Therefore feeding RNAi is robust in virtually all cells in *C. elegans* except neurons. To enhance neuronal RNAi, worms (TU3311 strain, uIs60[unc-119p::yfp ^+^ unc-119p::sid-1]) were used which express wild-type *sid-1* under control of the pan-neuronal promoter *unc-119* (Calixto et al. 2010). Synchronized L1 worms were transferred to NGM plates with RNAi-expressing bacteria. The control RNAi is the empty vector L4440. To prevent progeny, FUdR (200μM) was added before worms became adult. Online application for survival analysis (OASIS) was used to perform statistical analysis of the lifespan assays (Yang et al. 2011). To test whether RNAi inactivation of *F22D3.6* is accompanied by neurotoxicity, we performed a motility study with the WMicrotracker. This device is a high-throughput tracking system to record the amount of animal movement in a fixed period of time. The animal movement is detected through infrared microbeam light scattering. A 24-well plate, filled with nutrient agar, was used. Six wells were seeded with the control RNAi bacteria, six others were seeded with the RNAi bacteria against *F22D3.6*. Neuronal RNAi sensitive worms (TU3311 strain) were grown on RNAi bacteria until young adulthood (day 0). Around 100 adult worms were inoculated in each well. The exact number of the worms was counted afterwards. Three independent biological replicates were measured over a time period of 20 hours. Data acquisition and analysis was performed as described in (Simonetta et al. 2009). The detected signals per hour were divided by the average worm number in each well. The difference in motility was expressed relative to the control.

## Acknowledgments

Zebrafish cDNA was kindly provided by Prof. Kris Vleminckx (Ghent University). A pacific oyster (*Crassostrea gigas*) cDNA library was kindly provided by Prof. Pascal Favrel (UMR BOREA Biologie des ORganismes et Ecosystèmes Aquatiques, Institut de Biologie Fondamentale et Appliquée, Université de Caen Basse-Normandie). Work in the Beyaert lab is financed by the Fund for Scientific Research Flanders (FWO), the Belgian Foundation Against Cancer, Interuniversity Attraction Poles, Concerted Research Actions (GOA) and the Group-ID Multidisciplinary Research Partnership of Ghent University. A.B. is supported by a predoctoral fellowship from the FWO.

## Supplemental material

Supplemental text 1: FASTA sequences of type 1 paracaspases used in phylogeny

Supplemental text 2: FASTA sequences of Bcl10 homologs used in phylogeny

Supplemental text 3: FASTA sequences of CARMA/CARD9 homologs used in phylogeny

Supplemental text 4: FASTA sequences of Zap70/Syk homologs

**Supplemental Figure 1:**
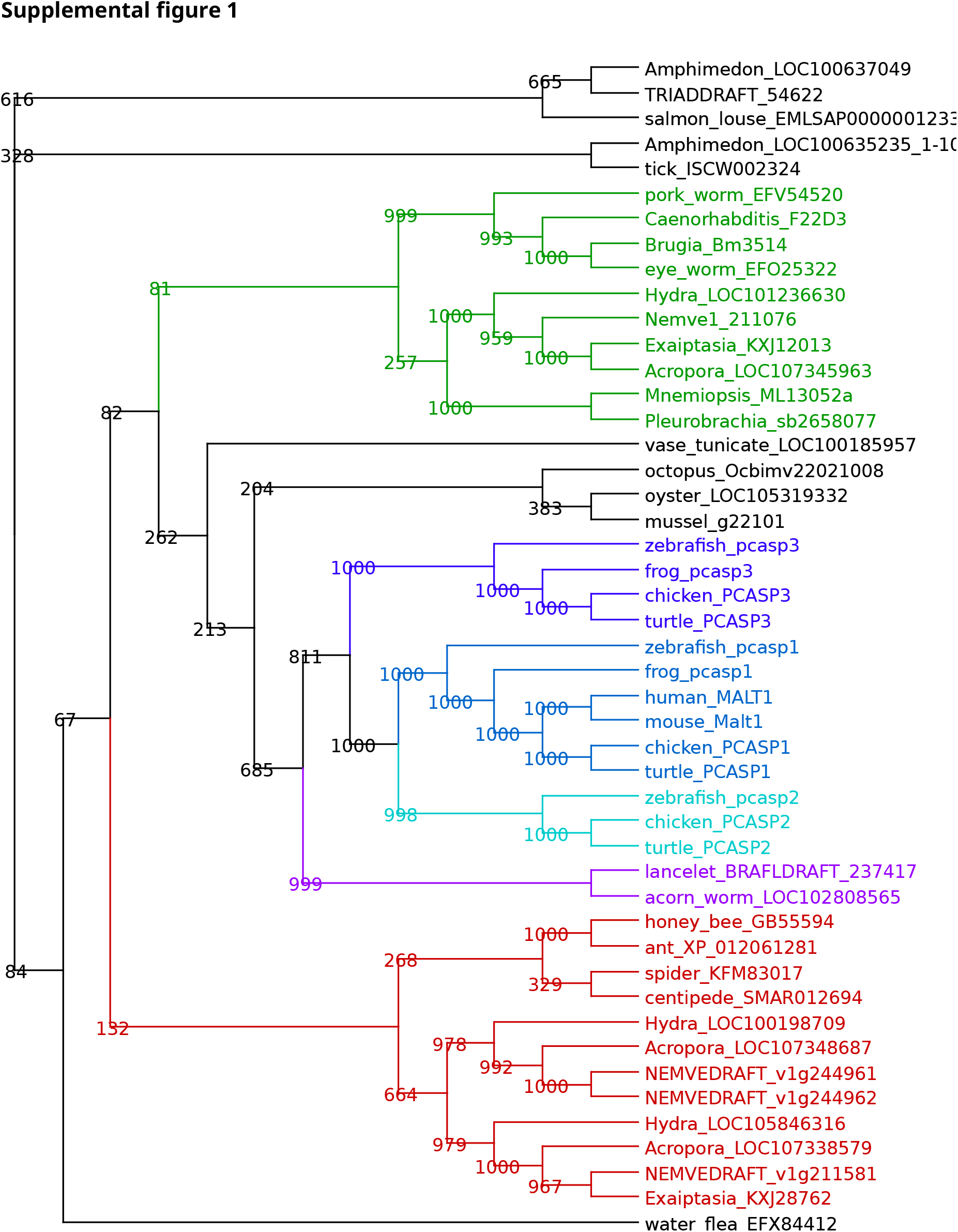
Detailed phylogenetic analysis of the type 1 paracaspases. Phylogenetic analysis of reliable full-length type 1 and type 2 paracaspases reveals 3 major families of type 1 paracaspases in bilaterians: The “arthropod” group (red), the “nematode” group (green) and the mollusk/deuterostome group (several colours). Arthropod and nematode type 1 paracaspases cluster with different type 1 paracaspases from cnidarians, indicating that the last common ancestor of planulozoans already had several type 1 paracaspase paralogs.

